# Predicting Isoniazid Resistance in *Mycobacterium tuberculosis* Complex in New York State using Whole Genome Sequencing

**DOI:** 10.1101/2025.10.30.685518

**Authors:** Kruthika Patel, Joseph Shea, Pascal Lapierre, Tanya A. Halse, Donna Kohlerschmidt, Michelle Dickinson, Vincent Escuyer, Kimberlee A. Musser

## Abstract

Isoniazid (INH) is a critical antibiotic used worldwide for treatment and prophylaxis of tuberculosis. Drug resistance (DR) to INH is the single most common type of DR, mediated by multiple genes/loci including *katG, inhA, mabA, mabA-inhA* and the *oxyR-ahpC* intergenic region. Over the course of 6 years, we performed a 2-phase study of 3,696 *Mycobacterium tuberculosis* complex (MTBC) strains aiming to determine the molecular basis of INH resistance and assess whole genome sequencing (WGS) for predicting resistance. In phase 1, we performed a side-by-side study including 1,767 strains with paired phenotypic drug susceptibility testing (DST) and genotypic DST. We found WGS capable of accurately predicting INH resistance with sensitivity of 90.3%, and specificity of 99.8%. The negative predictive value of WGS for INH susceptibility was 98.8%. Based on these findings, we developed a molecular testing algorithm where phenotypic DST was reduced and applied this new testing algorithm in phase 2 to 1,929 MTBC strains, resulting in streamlined testing, reduced cost and decreasing turnaround time (TAT). The prevalence of INH resistance among MTBC strains in New York was found to be 10.2%. Of the 3,696 isolates tested, 337 were predicted INH resistant by WGS. Of 41 additional strains exhibiting phenotypic INH resistance, 38 were found to have mutations in genes known to be associated with INH resistance. This study demonstrates the utility of WGS as a molecular tool for predicting INH DR and shows that the vast majority of INH resistance in MTBC has a molecular basis in known resistance loci.

## INTRODUCTION

Tuberculosis (TB) is one of the most prevalent infectious diseases caused by *Mycobacterium tuberculosis* complex (MTBC) bacteria. According to the World Health Organization (WHO), in 2023, an estimated 10.8 million people fell ill with TB worldwide and caused an estimated 1.25 million deaths in 2023, despite being a preventable and curable disease. TB continues to be the world’s leading cause of death from a single infectious agent (Global TB report, WHO, 2024) especially in people with acquired immune deficiency syndrome (AIDS) and a major contributor to the growing number of antimicrobial resistant infections, globally (Bastard et al., 2018, Gayoso et al., 2018; Kurbatova et al., 2012; Kliiman & Altraja, 2009). Multidrug-resistant TB (MDR-TB) is a form of TB caused by bacteria that are resistant to isoniazid (INH) and rifampin (RIF), the two most effective first-line TB drugs (Global TB report, WHO, 2014). According to the WHO guidelines, detection of MDR-TB requires bacteriological confirmation of TB and testing for drug resistance (DR) using rapid molecular tests, culture methods or sequencing technologies (Global TB report, WHO, 2022).

INH is one of the critical components of chemotherapy, used worldwide for both treatment and prophylaxis of TB. INH resistance often precedes RIF resistance in the development of MDR-TB (Manson et al., 2017). Therefore, rapid and accurate diagnosis of INH-resistant TB is essential to limit the spread of INH-resistant TB and reduce the likelihood of further acquisition of DR. Timely identification of MDR-TB cases will improve utilization of appropriate drug regimen treatments in patients and reduce the transmission of MDR-TB, resulting in better clinical outcomes (Gilpin et al., 2016). Mutations in several genes and intergenic regions including *katG*, *inhA*, *ndh*, *mabA (fabG1), mabA-inhA (fabG1-inhA),* and *oxyR-ahpC* have been associated with INH DR (Basso et al., 1998; Mdluli et al., 1998; Musser et al., 1996; Piatek et al., 2000; Ramaswamy et al., 1998, Sreevatsan et al., 1997). The *katG* locus encodes the catalase/peroxidase enzyme that is essential for INH activity as it activates the pro-drug INH to an active drug (Heym et al., 1993). Therefore, the presence of any mutation altering the activity of *katG* will prevent the activation of the pro-drug, resulting in resistance. Active INH disrupts the mycolic acid biosynthesis by inhibiting the product of *inhA*, the NADH-dependent enoyl acyl carrier protein (ACP) reductase enzyme (Unissa et al., 2016). Mutations in the promoter region cause the overexpression of INH’s target, NADH-dependent enoyl-ACP resulting in a MIC increase (Lempens et al., 2018).

Culture-based phenotypic drug susceptibility testing (DST) methods are still used frequently and remain the methods of reference for resistance profiling. However, these tests are slow and rely on mycobacterial growth in the presence of a critical drug concentration to distinguish between resistant and susceptible phenotype based on epidemiological breakpoints (Schon et al., 2017). Phenotypic DST is labor intensive, requiring specialized infrastructure and highly trained staff. Molecular methods to predict drug resistance can overcome some of the limitations of culture-based testing, particularly providing results at reduced turnaround time (TAT).

Molecular methods used to detect INH DR historically have included line probe assays (LPA); (Ling et al., 2008), Sanger sequencing of resistance-associated genes (Sekiguchi et al., 2007) and pyrosequencing (Halse et al., 2010). In 2016 the WHO approved the use of two LPAs: Hain GenoType MTBDR*plus* version 2 (Hain Lifescience, Nehren, Germany) and Nipro NTM + MDRTB detection kit 2 (Nipro, Tokyo, Japan). While these molecular assays can detect INH DR at greatly decreased TAT (Boehme et al., 2010), phenotypic DST is more sensitive at detecting resistance due to rare variants and resistance mechanisms which are not captured by these molecular tests. Newer methods such as next generation sequencing (NGS) have increased sensitivity of DR detection compared to previous molecular assays by increasing the number of resistance genes targeted. are important to expedite implementation of appropriate therapy and impact patients’ outcomes. NGS includes both targeted NGS (tNGS) and whole genome sequencing (WGS). tNGS assays are now endorsed by the WHO (Consolidated guidelines on TB, 2024) and include multiple commercially available tests as well as laboratory developed assays (Murphy et al., 2023).

Whole genome sequencing (WGS) is a powerful diagnostic tool, as it determines the complete DNA sequence of an organism’s genome but usually requires a cultured isolate. Unlike other molecular assays, WGS is capable of screening both known DR-associated genes and other less well-characterized loci in the MTBC genome. Therefore, a comprehensive list of mutations in target genes that confer DR can be generated, resulting in a reliable prediction of the drug susceptibility profile of a particular isolate, within a quicker timeframe.

The aims of this study were to determine the capacity of WGS to predict INH resistance and susceptibility of clinical MTBC isolates in New York State (NYS), and to evaluate a testing algorithm with reduced phenotypic DST. In the first phase of this study, paired phenotypic and genotypic data were compared to determine the molecular basis of resistance in strains exhibiting phenotypic INH resistance. These data were used to inform the construction of a testing algorithm with WGS as the primary method of susceptibility profiling and phenotypic DST as a reflex method for non pan-susceptible strains, which was implemented for the second phase of this study. We found WGS capable of predicting INH resistance and susceptibility with high sensitivity and specificity, which led us to use WGS as our primary method of drug susceptibility testing. Strains with rare and novel mutations continue to have phenotypic testing performed to contribute to the growing knowledge base of INH resistance mechanisms and mutations and to improve the predictive power of molecular assays.

## MATERIALS AND METHODS

### Clinical isolates

A total of 3,696 MTBC strains were included in this study consisting of one isolate from every culture-positive case in NYS over a period of six years between January of 2016 and February of 2022. All samples were received as isolates or cultured in-house from clinical specimens in the biosafety level 3 (BSL-3) Mycobacteriology Laboratory at the Wadsworth Center, New York State Department of Health (NYSDOH). An in-house developed real-time PCR assay was used to confirm the presence of MTBC DNA in all the specimens and isolates that were received (Halse et al., 2010).

### DNA extraction

DNA extraction was performed on heat inactivated (80°C for 1 hour) liquid culture aliquots of clinical isolates using a modified version of the InstaGene/FastPrep (IG/FP) method (Shea et al., 2017). Specifically, the volume of InstaGene added to the pellets was changed to be 130 – 200 µL based on the size of the pellet to increase DNA yield, and the 56°C incubation was reduced from 30 min to 10 min.

### Phenotypic drug susceptibility testing

Phenotypic DST was performed using the liquid culture MGIT 960 system ((BACTEC Mycobacterial Growth indicator tube (MGIT) 960 SIRE package insert; Becton, Dickinson) and solid 7H10 agar proportion method according to the Clinical and Laboratory Standards Institute’s recommendations (CLSI M24 3^rd^ edition). Prior to DST being set up, isolates were sub-cultured in MGIT medium. MIC testing on a subset of isolates was performed by broth microdilution using Sensititre plates (ThermoFisher) to further characterize mutations associated with INH resistance. The range of concentrations tested for INH was 0.025 to 12.8 µg/mL. MIC testing was performed in triplicate, with plates being read by one analyst.

### Whole genome sequencing

Paired-end sequencing was performed on the Illumina MiSeq® and Nextseq® platforms following Nextera Flex and Illumina DNA Prep library preparation, respectively. Sequencing runs were either all MTBC isolates or MTBC isolates with other bacterial, viral and/or parasitic samples as previously described (Shea et al., 2017) but with improvements in cost and turn-around time described in Dickinson et al (2024).

### Bioinformatic analysis

Sequence analysis was performed using the Wadsworth Center TB WGS bioinformatics pipeline as previously described to detect an internally curated list of high confidence (HC) genomic insertions and deletions (indels) and single nucleotide polymorphisms (SNPs) for drug resistance profiling (Shea et al., 2017). Furthermore, all mutations in INH associated genes and gene regions were screened to identify rare and novel resistance mutations. Major lineage identification was based on the presence or absence of lineage defining SNPs, with nomenclature according to Gagneux and Small (2007) and Feuerriegel et al. (2014).

## RESULTS

Universal WGS was performed on one isolate from every TB case in NYS with an available MTBC isolate over a six-year period (n = 3,696). In the first phase of this study (01/2016 – 09/2018) side-by-side phenotypic DST (MGIT & agar proportion) and genotypic testing (WGS) was performed for all samples (n = 1,767). In the second phase (10/2018 – 02/2022), a tiered testing algorithm was implemented (n=1,929) where phenotypic DST was limited to samples with known DR mutations (HC mutations) and mutations of unknown significance, to reduce the testing burden while still collecting phenotypic DST data to link with rare mutations and mutations of unknown significance (Shea et al., 2023).

### Study Phase 1

WGS showed high levels of concordance with phenotypic DST (90.3% sensitivity & 99.8% specificity, Table 1 & 2). Among the 1,767 isolates, 169 were identified as resistant and 1,598 as susceptible by WGS. Among the isolates with no high confidence mutations detected by WGS, 14 isolates exhibited phenotypic DR. Of these 13 harbored one or more mutations in genes associated with INH resistance (10 *katG*, 1 *katG* + *oxyR-ahpC*, 1 *inhA,* and 1 *oxyR-ahpC + furA-katG*). Of the 169 isolates predicted resistant by WGS, phenotypic DST confirmed 130 as resistant and reported two as susceptible (*mabA-inhA*: T-8A and *katG*: insertion). The remaining 37 had no phenotypic DST data available however had well-known mutations in *katG* and *mabA-inhA* intergenic region which are associated with resistance to INH (Seifert et al., 2015, Vilcheze et al., 2014). Among the INH-resistant isolates (n=169), 157 had a single HC mutation and 12 had two HC mutations. (Table 3).

**Table 1:**
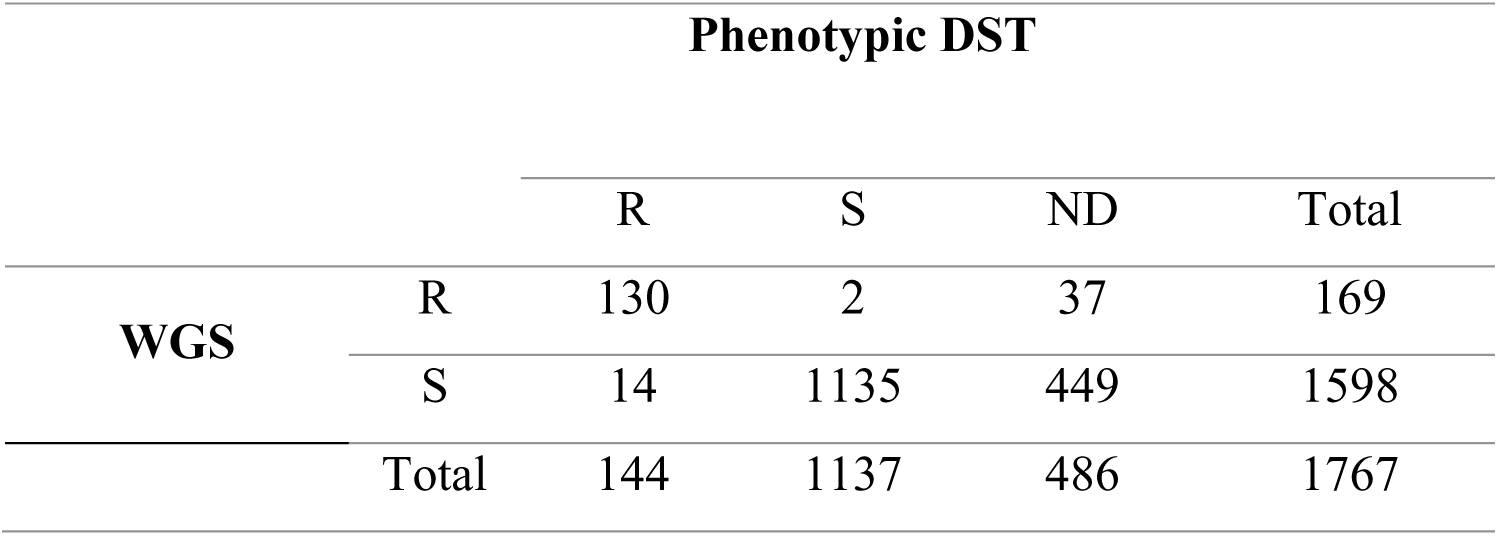
Summary Statistics of WGS Determined High-Confidence Mutations and pDST During Phase 1.

**Table 2:** Summary Statistics of WGS High-Confidence Mutations with pDST and INH Resistance Prevalence During Phase 1.

| Performance characteristics | Phase 1 |
| --- | --- |
| Sensitivity % (n) | 90.3 (130/144) |
| Specificity % (n) | 99.8 (1135/1137) |
| Positive (resistance) predictive value % (n) | 98.5 (130/132) |
| Negative (susceptibility) predictive value % (n) | 98.8 (1135/1149) |
| Accuracy % (n) | 98.7 (1265/1281) |

Phase 1: 1767 total isolates, 486 samples did not have pDST data available.

**Table 3:** Frequency of HC Mutations in 169 INH-Resistant Isolates During Phase 1.

| DST pattern | N | Amino Acid Change |  |  |  |  |
| --- | --- | --- | --- | --- | --- | --- |
|  |  | *katG* | *inhA* | *mabA* | *mabA-inhA* | *oxyR-ahpC* |
| INH Resistant | 64 | Ser315Thr |  |  |  |  |
|  | 3 | Ser315Asn |  |  |  |  |
|  | 3 | Deletion |  |  |  |  |
|  | 1 | Insertion |  |  |  |  |
|  | 2 |  | Ser94Ala |  |  |  |
|  | 8 |  |  | Leu203Leu |  |  |
|  | 34 |  |  |  | C-15T |  |
|  | 3 |  |  |  | G-17T |  |
|  | 3 | Ser315Thr |  | Leu203Leu |  |  |
|  | 3 | Ser315Thr |  |  | C-15T |  |
|  | 1 | Ser315Thr |  |  | T-8C |  |
|  | 2 | Gly121Ser |  |  | C-15T |  |
|  | 1 | Gln525Pro |  |  | C-15T |  |
|  | 1 | Deletion |  |  |  | C-81T |
|  | 1 |  | Ser94Ala |  | C-15T |  |
| INH Susceptible | 1 |  |  |  | T-8A |  |
|  | 1 | Insertion |  |  |  |  |
| INH Not Determined | 26 | Ser315Thr |  |  |  |  |
|  | 3 | Deletion |  |  |  |  |
|  | 8 |  |  |  | C-15T |  |
| Total | 169 |  |  |  |  |  |

### Study Phase 2

A WGS prediction scheme categorizing MTBC strains as INH-resistant (HC mutations), unknown, or susceptible (R/U/S) based on the mutation profile was implemented and strains predicted to be susceptible had no phenotypic DST performed. The prediction category of unknown allows for targeted phenotypic testing to be performed on a subset of samples that are more likely to be INH resistant. Mutations of unknown significance which were found to have resistant phenotype were most commonly found in *katG*. During this phase of the study, the list of HC mutations used to predict INH resistance was updated to include *inhA* Ser94Ala, *katG* Gly121Asp, Trp191Gly and Thr394Ala.

Among the 1,929 isolates, 168 were predicted as INH resistant, 1,651 as INH susceptible and 111 as unknown by WGS. Of the 111 with mutations of unknown significance, 25 exhibited phenotypic resistance, 15 of which had mutations in *katG* gene, six had mutations in *katG* and *oxyR-ahpC*, two had mutations in the *furA-katG* promoter region, and one had a mutation in *mabA*. Among the isolates predicted susceptible by WGS, two had no mutations in known resistance loci, and no explanation for the phenotypic resistance. Of the 168 predicted resistant by WGS, phenotypic DST confirmed 97 as resistant. The remaining 71 had no phenotypic DST data available however had well-known mutations in *katG, inhA, mabA* and *mabA-inhA* promoter region which are associated with INH resistance (Seifert et al., 2015, Vilcheze et al., 2014). Among the INH-resistant isolates (n=168), 163 had a single HC mutation and five had two HC mutations (Table 4).

**Table 4:**
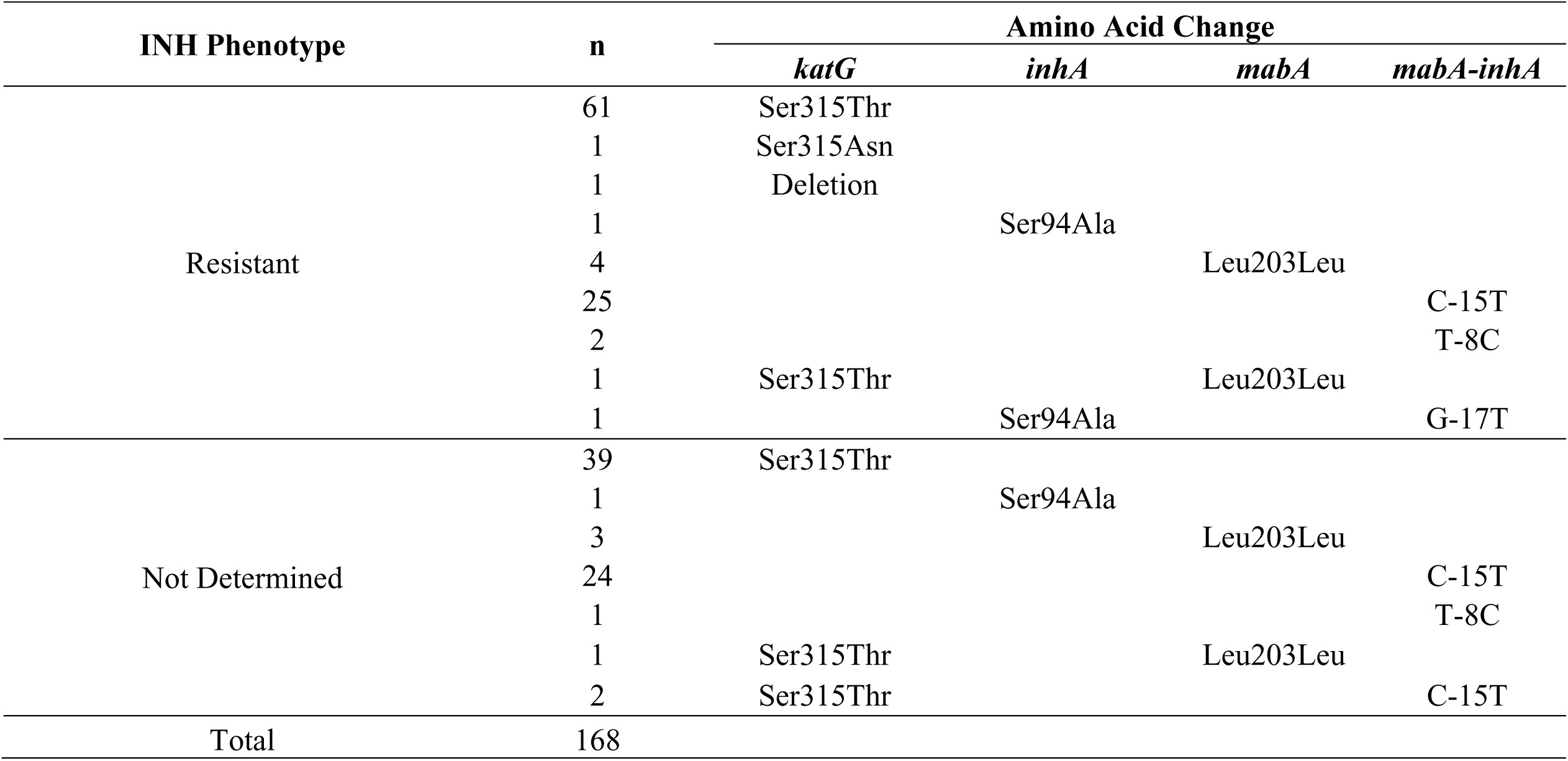
Frequency of HC mutations in 168 INH-Resistant Isolates During Phase 2.

| INH Phenotype | n | Amino Acid Change |  |  |  |
| --- | --- | --- | --- | --- | --- |
|  |  | *katG* | *inhA* | *mabA* | *mabA-inhA* |
| Resistant | 61 | Ser315Thr |  |  |  |
|  | 1 | Ser315Asn |  |  |  |
|  | 1 | Deletion |  |  |  |
|  | 1 |  | Ser94Ala |  |  |
|  | 4 |  |  | Leu203Leu |  |
|  | 25 |  |  |  | C-15T |
|  | 2 |  |  |  | T-8C |
|  | 1 | Ser315Thr |  | Leu203Leu |  |
|  | 1 |  | Ser94Ala |  | G-17T |
| Not Determined | 39 | Ser315Thr |  |  |  |
|  | 1 |  | Ser94Ala |  |  |
|  | 3 |  |  | Leu203Leu |  |
|  | 24 |  |  |  | C-15T |
|  | 1 |  |  |  | T-8C |
|  | 1 | Ser315Thr |  | Leu203Leu |  |
|  | 2 | Ser315Thr |  |  | C-15T |
| Total | 168 |  |  |  |  |

### Combined datasets summary

Of a total of 3,696 isolates tested during phase 1 and 2, 337 were found to have HC mutation(s) in one or more genes associated with INH resistance (*katG, mabA-inhA, mabA, inhA, oxyR-ahpC*) (Table 5). Of these, 312 (92.6%) isolates had a mutation in a single gene/locus, while 16 (4.7%) had mutations in multiple loci. The remaining nine (2.7%) had indels in *katG*. Of the 337 INH resistant isolates, 60.2% of HC mutations were found in *katG* (n=203) followed by *mabA-inhA* (n=98; 29.1%), *mabA* (n=15; 4.4%), *inhA* (n=4; 1.2%), oxyR-ahpC (n=1; 0.3%) and the remaining 4.7% had HC mutations in two separate genes (Figure 1). The predominant HC mutation was *katG* Ser315Thr, which was found in 190/337 (56.4%) of the strains predicted INH resistant by WGS. Other frequent HC mutations included *mabA-inhA* C-15T in 91 strains (27.0%) and *mabA* Leu203Leu (CTG to CTA) in 15 strains (4.5%). Among 268 total isolates found to have phenotypic INH resistance, 227 (84.7%) had HC mutations and 39 (14.5%) had mutations of unknown significance in genes associated with INH resistance. In total, 266/268 (98.9%) strains with phenotypic INH resistance were found to have a possible molecular basis of resistance by WGS.

**Figure 1:**
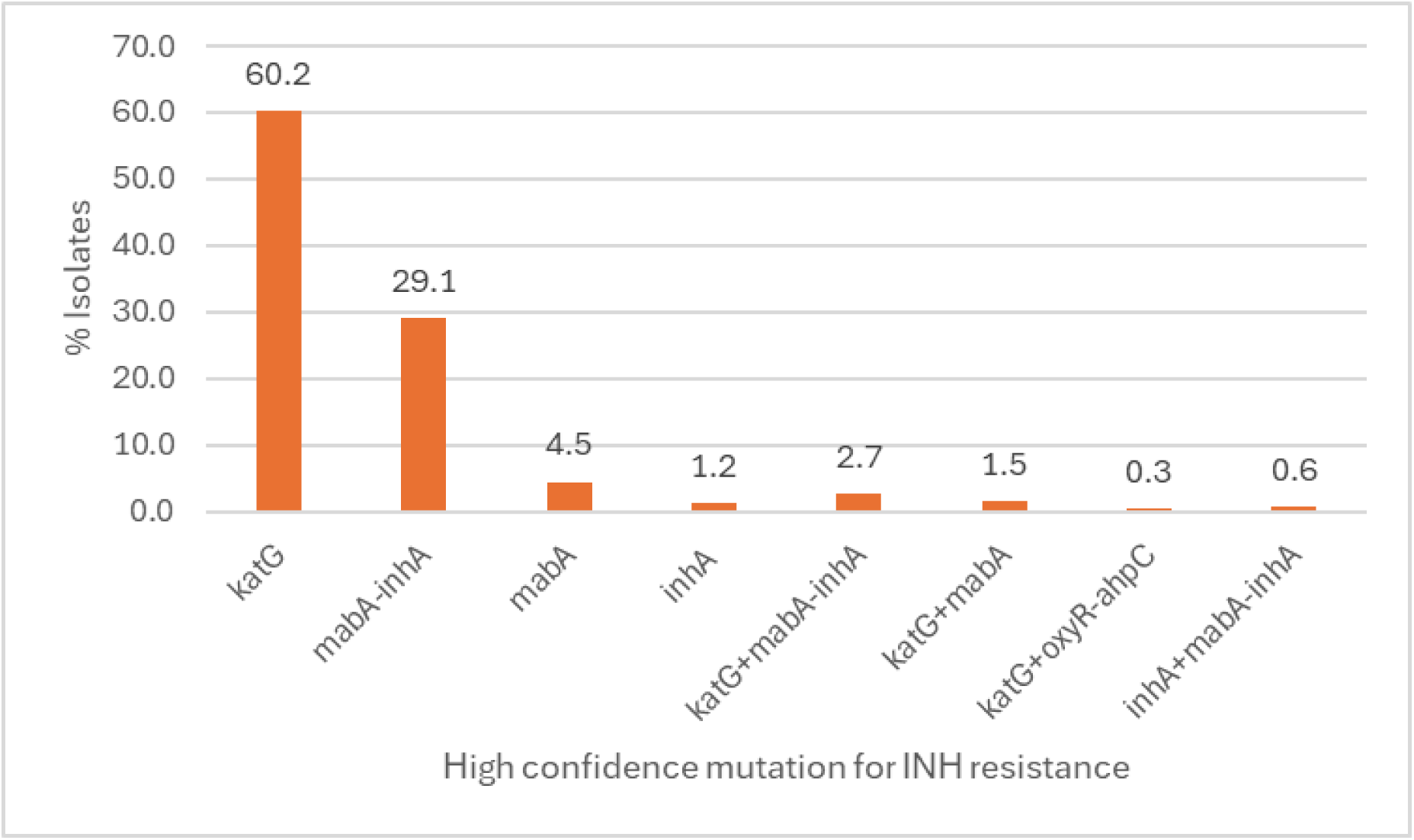
Proportion of resistance mutations in each gene/gene region in INH-resistant MTBC isolates (n=337)

**Table 5:** WGS Predictions and Corresponding pDST Results for INH in 3696 MTBC Isolates.

|  | pDST Resistant | pDST Susceptible | pDST Not determined | Total |
| --- | --- | --- | --- | --- |
| **WGS Resistant** | 227 | 2 | 108 | 337 |
| **WGS Susceptible** | 17 | 1329 | 1903 | 3249 |
| **WGS Unknown** | 24 | 78 | 8 | 110 |
| **Total** | 268 | 1409 | 2019 | 3696 |

### Phenotypic drug susceptibility testing

DST data were available for 1,677 (45.4%) MTBC isolates based on MGIT DST. Of these, 268 (15.9%) were INH resistant with 227 harboring HC mutations. The remaining 41 isolates exhibited phenotypic resistance of which 39 harbored mutations of unknown significance in resistance conferring genes (Table 6). MIC data was available only on a subset of the isolates included in this study. Of the 337 INH-resistant isolates, 75 (22.3%) had MIC data available and the distribution of corresponding HC mutations with level of resistance is shown in Figure 2. When tested by broth microdilution assay, the majority of the strains (50/75) were highly resistant to INH with MIC values over 2 µg/mL and had mutations in the *katG* gene codon 315. Of these, 30/50 had an MIC ≥4 µg/mL. Mutations in other genes explored in this study conferred MIC values ranging from 0.12 to 1 µg/mL. For example, the most common mutation in the mabA-inhA intergenic region, C-15T had an MIC ranging from 0.25 to 1 µg/mL and the less common mutations such as T-8A and G-17T had an MIC of 0.12 and 0.25 µg/mL, respectively. The silent mutation Leu203Leu in the mabA gene had an MIC of 0.12 – 0.25 µg/mL.

**Figure 2:**
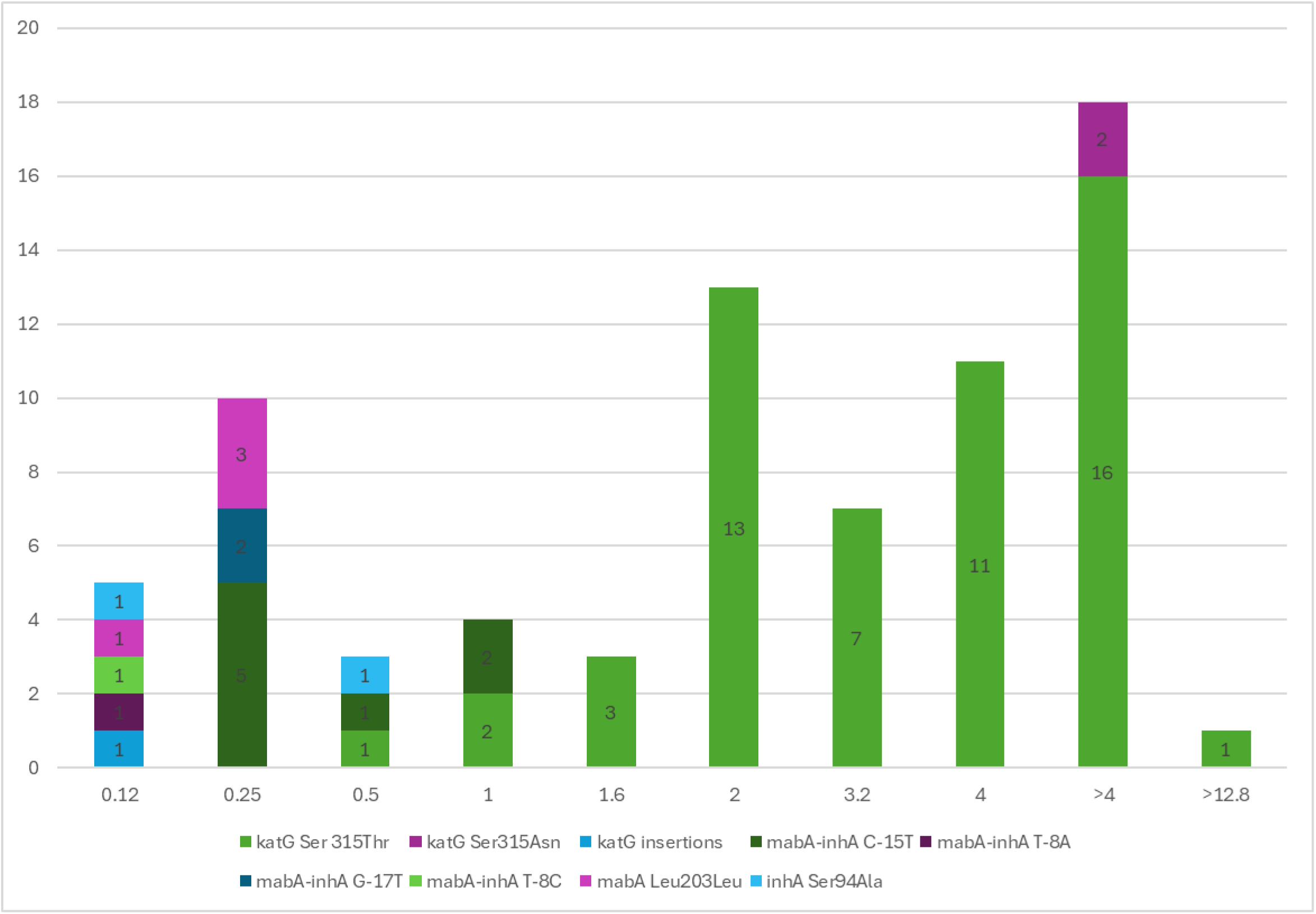
INH MIC distribution of HC mutations by genes in MTBC isolates (n=75)

**Table 6:** Potential Resistance Mutations in Isolates that were Phenotypically INH-Resistant (n = 39)

| Resistance Gene/ Gene Region | Amino Acid/Nucleotide Change | No. of Isolates |
| --- | --- | --- |
| *katG* | Ala93Val | 1 |
|  | Ala144Val | 1 |
|  | Ala162Thr | 1 |
|  | Arg128Trp | 1 |
|  | Arg249Gly | 1 |
|  | Arg484Cys | 1 |
|  | Arg595Gln | 3 |
|  | Asp94Ala | 1 |
|  | Gly121Asp | 1 |
|  | Gly123Glu, Ala706Glu | 1 |
|  | Leu427Pro | 1 |
|  | Phe408Val | 1 |
|  | Phe540Ser | 1 |
|  | Ser315Gly | 1 |
|  | Thr271Pro | 1 |
|  | Thr344Pro, Gln352His | 1 |
|  | Thr394Ala | 2 |
|  | Trp191Gly | 1 |
|  | Trp300Arg | 1 |
|  | Tyr597Cys | 1 |
| *katG + oxyR-ahpC* | Ala109Thr + T-77A | 1 |
|  | Gln127Pro, Lys157Glu + C-52A | 1 |
|  | Gly299Ala + C-54T | 1 |
|  | Met176Val + C-72T | 1 |
|  | Trp191Gly + G-48A | 1 |
|  | Trp191Arg + G-51A | 1 |
|  | Val731Gly + C-52T | 1 |
| *furA-katG* | A-9C | 2 |
| *katG + ndh* | Val1Val (GTG to GTA) + Gly313Arg | 1 |
| *furA-katG + oxyR-ahpC* | A-10C + C-54T | 1 |
| *inhA* | Ile21Val | 1 |
| *ndh* | Arg268His | 1 |
|  | Arg284Trp | 1 |
|  | Insertion | 1 |
| *mabA* | Val221Val (GTC to GTA) | 1 |

### INH resistance and mutation prevalence by major lineage

All four major lineages of *M. tuberculosis* (L1, L2, L3 and L4) were represented in our data set (14.4%, 22.3%, 8.4%, and 51.2% of strains, respectively). The distribution of strains with INH resistance-conferring mutations across these four lineages varied drastically, ranging from 19.3% in L1, 32.6% in L2, 3.9% in L3 and 43.9% in L4. INH resistance mutations were over-represented in L1 and L2 strains, given the prevalence of these lineages in this study (Table 7). We also found 131 strains to be other members of the MTBC (52 *Mycobacterium bovis*-BCG, 37 *Mycobacterium bovis*, 34 *Mycobacterium africanum*, 6 *Mycobacterium orygis*, 1 *Mycobacterium caprae*, and 1 strain of novel lineage La4 (Shea et al., 2022); none of these strains exhibited any INH resistance.

**Table 7:** Isoniazid Resistance and Mutation Prevalence by Major Lineages.

| Lineage | No. of Strains (%) | No. of INH-Resistant Strains by WGS (%) |
| --- | --- | --- |
| L1 | 531 (14.4) | 65 (19.3) |
| L2 | 526 (22.3) | 110 (32.6) |
| L3 | 312 (8.4) | 13 (3.9) |
| L4 | 1892 (51.2) | 148 (43.9) |
| *M. bovis* BCG | 52 (0.01) |  |
| *M. bovis* | 37 (0.01) |  |
| *M. africanum* | 34 (0.009) |  |
| *M. orygis* | 6 (0.002) |  |
| L7 | 1 (0.0003) |  |
| La4 | 1 (0.0003) |  |
| Mix of 2 Lineages | 1 (0.0003) | 1 (0.003) |
| *M. caprae* | 1 (0.0003) |  |
| **Total** | 3696 | 337 |

## DISCUSSION

The last decade has seen tremendous advances in NGS technologies including the use of WGS as part of the routine clinical diagnosis for infectious diseases. Since implementation in our laboratory in January of 2016, clinical WGS for MTBC has improved surveillance and detection of INH resistance while providing insights into the molecular basis of INH-resistant strains in NYS. In this six-year study, we analyzed a set of 3,696 MTBC isolates from unique patients to assess the frequency and type of INH resistance-conferring mutations determined by WGS and their association with phenotypic DST test results, including MIC testing. Additional data were collected to evaluate the relationship of strain lineage with INH resistance.

Past reports have shown that mechanisms of INH resistance are highly complex due to the involvement of several genes and intergenic regions (Unissa et al., 2016). In this study we looked at the mutations in three genes and two regulatory regions associated with INH resistance. The predominant loci in which mutations were associated with INH resistance were *katG* and *mabA-inhA*, with a smaller proportion of INH-resistance associated with *mabA*, *inhA* and *oxyR-ahpC* mutations.

Mutations in *katG* confer INH resistance by blocking the pro-drug activation pathway. A variety of resistance conferring mutations have been identified thus far in *katG*, though mutations in codon 315 are the most frequent and often confer moderate to high levels of INH DR (Dantes et al., 2012). In the current study, the prevalence of *katG* mutations (60.2%) conferring INH resistance was consistent with previous reports which have shown that mutations in the *katG* gene are the most prevalent, with 60-67% of INH-resistant clinical TB isolates harboring mutations in this locus (Lempens et al., 2018; Tseng et al., 2013; Jabbar et al., 2019, Seifert et al., 2015). Among the INH-resistant isolates with *katG* codon 315 mutations, the most common nucleotide change at *katG* codon 315, AGC to ACC (Serine to Threonine), occurred in 97.9% of the isolates. This is similar to a systematic review published in 2015, which reported that 93.4% of the phenotypic INH resistance was associated exclusively with a single Ser315Thr mutation in *katG* (Seifert et al., 2015). Despite this mutation, the catalase/peroxidase activity of the enzyme is maintained but has reduced ability to activate INH (Gygli et al., 2017; Torres et al., 2015). The Ser315Asn mutation is also an INH resistance-conferring mutation that occurs at a lower frequency. Its prevalence in the present study, 4 out of 194 isolates (2.0%) with mutations in the *katG* codon 315, is closer to that of the global average (3.6%) reported by Seifert et al, 2015. Mutations in the *mabA-inhA* intergenic region confer INH resistance by overexpression of the drug target resulting in removal of the drug from the bacterial cell. Such mutations have been associated with a lower level of phenotypic DR compared to other mutations such as the common *katG* Ser315Thr (Lempens et al., 2018). In the current study, 29.1% of the INH resistant isolates had mutations in *mabA-inhA* gene which is consistent with other studies that have reported 20-42% of clinical MTBC isolates with INH resistance having mutations in this intergenic region (Bakhtiyariniya et al., 2022; Rattan et al., 1998; Rossetti et al., 2002; Ahmad et al., 2009; Sreevatsan et al 1997, Seifert et al., 2015).

The most prevalent *mabA-inhA* intergenic region mutation is the C-15T mutation (Seifert et al., 2015) present in 92.2% of INH-resistant isolates with mutation in this intergenic region worldwide which is similar to what was observed in this study (92.8%, 91 of 98). Although C-15T mutation is the dominant mutation in the *mabA-inhA* intergenic region, other resistance-associated mutations (7%) were observed and occur independently of the C-15T mutation. Combinations of mutations in *katG* and the *mabA-inhA* intergenic region are known to confer high-level resistance (Kambli et al., 2015). In the current study, five isolates carrying C-15T mutation also had the Ser315Thr mutation in the *katG* gene.

A synonymous mutation in *mabA*, Leu203Leu (CTG to CTA), confers INH resistance by creating an alternative promoter for *inhA*, resulting in its overexpression (Bainomugisa et. al., 2022). In most cases silent mutations do not play a role in DR, however, the Leu203Leu mutation demonstrates that a silent mutation can be associated with DR depending on the specific gene and the location of the mutation. Another locus for mutations in INH-resistant MTBC isolates is the *oxyR-ahpC* regulatory region. Mutations in this region of *katG*-defective INH-resistant MTBC strains have been described which result in upregulation of AhpC, an alkyl hydroperoxidase, and thus compensate for the loss of catalase-peroxidase activity (Kelly et al., 1997; Ramaswamy et al., 2003; Clemente et al., 2008; Sreevatsan et al., 1997). However, mutations in the *oxyR-ahpC* regulatory region have also been identified in INH-susceptible MTBC isolates, suggesting that these mutations themselves may not results in INH resistance, but may be selected for in strains which are INH resistant (Hazbon et al., 2006). Nine INH resistant isolates were found to have mutations in *oxyR-ahpC*, numbered in this study based on position relative to the start codon of *ahpC*, (G-48A, G-51A, C-52A, C-52T, two C-54T, C-72T, T-77A, C-81T) however each of these had a mutations in *katG* which could account for the phenotypic resistance. These mutations are believed to result in AhpC overexpression and partially restore the fitness cost due to reduced *katG* activity (Gygli et al., 2017).

The MIC for *Mycobacterium tuberculosis* to INH varies significantly, with sensitive strains typically having MICs below 0.2 μg/ml, while isoniazid resistance is associated with higher MICs, ranging from above 2 μg/ml up to >10 μg/ml. These resistance levels are often linked to mutations in the *<u>katG</u>*or *<u>inhA</u>* genes, though the specific MIC values can overlap and are influenced by the particular mutation present (Ramaswamy et al., 2003; Jagielski et.al., 2013, Kambli et al., 2015). Our results are largely in agreement with the associations between known resistance-conferring mutations and the level of phenotypic resistance described in the literature. MICs for INH-resistant isolates with Ser315Thr mutation in *katG* (n=54), were in the range of 0.5 to ≥12.8 µg/mL, and the MICs for isolates with amino acid replacements other than threonine at codon 315 of *katG* were ≥4 µg/mL (Figure 2). Previous studies have shown that 61.7%–100% of the mutations at codon 315 in the *katG* gene, most commonly Ser315Thr, exhibit higher levels of INH resistance (Lempens et al., 2018, Dantes et al., 2012, Abe et al., 2008, van Soolingen et al., 2000).

In our analysis, isolates with mutations in the *mabA-inhA* intergenic region (n=12) had MIC values ranging from 0.12 to 1 µg/mL, corroborating the findings by Lempens et al., who reported that 50% of the isolates with C-15T mutation showed lower levels of INH resistance. The same study also reported that 5.8% of the isolates with C-15T mutation had higher MIC values ranging from 3.2 to 12.8 µg/mL. Higher level INH resistance in isolates carrying C-15T mutation can be explained if it coincides with a mutation in *katG*, especially Ser315Thr (Lempens et al., 2018, Liu et al., 2018) or by mutations in other resistance loci.

We provide further evidence that much of the resistance to INH in MTBC can be explained by mutations in known resistance loci. Of the 268 isolates with phenotypic INH resistance 98.9% had mutations which could account for the phenotypic resistance. While most of these had HC mutations (84.7%), the remaining 38 isolates had less well characterized mutations and required phenotypic testing for final determinations of resistance. Five of the 39 isolates had *katG* mutations (One Gly121Asp, Two Thr394Ala, Two Trp191Gly) that we confirmed to be associated with INH resistance and have now been included in our in-house pipeline as HC mutations. Of the remaining 33 isolates, 10 had *katG* mutations that are mentioned in the WHO catalogue as mutations of uncertain significance and 2 had *ndh* mutations associated with intermediate INH resistance (WHO catalogue), however recent studies have shown that *ndh* is not involved in INH resistance (Pandey et al., 2024). Targeted testing to identify only the most common resistance mutations such as *katG* Ser315Thr and *inhA* C–15T cannot fully replace conventional DST, and this study provides further evidence that less common mutations in *katG, inhA, mabA,* and *mabA-inhA* also account for INH resistance in clinical strains. Early detection of these less common INH resistance-associated mutations could decrease the chances of treatment failure and additional acquisition of DR.

This study explored the prevalence of INH resistance-associated mutations in three genes and three regulatory gene regions reported to be associated with INH resistance in 3,696 clinical MTBC isolates. Based on the strong correlation observed between WGS and phenotypic DST in phase 1, a WGS prediction scheme categorizing the presence or absence of mutations as predictive of INH resistant, unknown, or susceptible (R/U/S) phenotypes and a tiered testing algorithm was implemented in phase 2. In phase 2 phenotyping testing was limited to samples with known resistance mutations and mutations of unknown significance. Some of the mutations of unknown significance were critical to detect instances of INH resistance which would have otherwise gone undetected; this was primarily due to rare mutations in the *katG* gene. Our findings in this study highlight the importance of tracking mutations of unknown significance and linking with phenotypic testing results to improve molecular predictions of INH resistance. This work will continue to contribute in the long-term to improvement of our WGS prediction algorithm for INH DR.

WGS is a valuable tool for identifying genome-wide variants and proves to be effective in detecting resistance to INH. Implementation of TB WGS has resulted in the rapid and accurate identification of INH resistance and multi-drug-resistance in MTBC isolates in NYS. In the context of a reduced phenotypic DST testing algorithm, WGS has also decreased the amount of phenotypic testing required while still reliably detecting INH resistant tuberculosis. Over time this work contributes to the efforts to characterize the molecular basis of INH resistance in tuberculosis and to improve the power of molecular assays at predicting drug resistance.

## ACKNOWLEDGEMENTS

This research project was partially supported by Cooperative Agreement #1U60OE000103 (CFDA NO. 93.322) with the Association of Public Health Laboratories and the U.S. Centers for Disease Control and Prevention (CDC). We also want to acknowledge the Wadsworth Center Advanced Genomic Technologies and Bioinformatics cores for their support for this testing.

